# Ramped V1 transcranial ultrasonic stimulation modulates but does not evoke visual evoked potentials

**DOI:** 10.1101/2023.01.24.525317

**Authors:** Tulika Nandi, Ainslie Johnstone, Eleanor Martin, Robert Cooper, Sven Bestmann, Til Ole Bergmann, Bradley Treeby, Charlotte J. Stagg

**Author notes:** **Corresponding author –** Tulika Nandi, NeuroImaging Center (NIC), Johannes Gutenberg University Medical Center, Langenbeckstr. 1, Bldg. 308c, 55131 Mainz.

## Abstract

Transcranial ultrasonic stimulation (TUS), has been shown to evoke ‘visual evoked potential (VEP)-like’ potentials on EEG recordings, and also to modulate sensory evoked potentials. However, pulsed TUS is accompanied by an auditory confound, and it is possible that any observed effects were, in-part, evoked by this confound. Therefore, we used ramped TUS pulses, which are not as easily audible, to examine whether primary visual cortex (V1) TUS evokes VEP-like potentials, and modulates VEPs elicited using a checkerboard stimulus.

**Methods:** We tested 14 healthy participants (31 ± 4.3 yrs, 4 F and 10 M). TUS was applied to the left V1 using a 270 kHz transducer (H115-2AA, Sonic Concepts). Ramped pulses (1 ms ramp, 3.25 ms total pulse duration) were repeated at 250 Hz, with a pulse train duration of 300 ms, an effective duty cycle of 50%, and Isppa without ramping of 16 W/cm^2^ in water. EEG was recorded from 16 channels using the g.USBamp amplifier (g.tec medical engineering GmbH). In two blocks (TUS-only), real and sham (100 each) TUS trials were repeated every 2 s. In another two blocks (TUS+checkerboard), a checkerboard stimulus was flipped every 0.5 s, and every fourth stimulus was associated with either a real or sham (100 each) TUS trial. The TUS trial started approx. 130 ms (0-5 ms jitter) before the checkerboard flip. All EEG data were analysed using Fieldtrip, and cluster-based permutation tests were used to test for differences between conditions.

**Results and discussion:** In the TUS-only condition, in contrast to a previous study, we found no evoked potentials using ramped pulses which minimised the auditory artifact. In the TUS+checkerboard condition, we observed a modulation of the early-component of the VEP in real TUS, relative to no TUS trials. This suggests that, in line with in-vitro and animal data, there is a direct neuromodulatory effect of ultrasound, in addition to any confounding effects. Moving forward, ramping offers a relatively easy approach to minimise the auditory confound.

---

Transcranial ultrasonic stimulation (TUS) is an emerging non-invasive brain stimulation technique that has higher spatial resolution than electrical and magnetic stimulation approaches, and, uniquely, offers the ability to target structures deep in the brain. Early work in humans suggests that TUS can both evoke neural activity (1) and modulate activity elicited by other stimuli (2). However, the protocols used in these studies may be audible due to the sharp onset and offset of ultrasound energy (3,4), and it is therefore possible that there is an auditory confound to the observed effects (5). Here, therefore, we used a less audible, ramped protocol (3) to determine if we could either evoke or modulate activity in the primary visual cortex (V1).

We examined whether V1 TUS alone evokes neural activity detectable in the EEG, and whether TUS modulates visual evoked potentials (VEPs) in response to a pattern-reversal checkerboard stimulus. Fourteen healthy participants (4 female, 31 ± 4.3 years) were included in the study, after excluding three participants due to technical problems. The project was approved by the UCL research ethics committee (Project ID 14071/001).

As described in our previous paper (3), TUS was delivered using a 2-element spherically focusing annular array transducer (H115-2AA, Sonic Concepts) with a nominal outer aperture diameter and radius of curvature of 64mm. The transducer was driven at 270 kHz by a 2-channel TPO (Sonic Concepts) with the output power and element phase adjusted to give a focal pressure in water of 700 kPa (spatial peak pulse average intensity without ramping of 16 W/cm^2^) and a focal distance of 43 mm. The measured -3 dB focal size in water was 5 mm (lateral) by 30 mm (axial). Ramped pulses (1 ms Tukey ramp, 3.25 ms total pulse duration) were applied at a pulse repetition frequency (PRF) of 250 Hz, with a pulse train duration of 300 ms and an effective duty cycle of 50%. Of the 14 participants, 7 could not hear the stimulation at all, while the rest were either uncertain or heard it faintly. EEG was recorded from 16 channels (Fig. 1a), at 600 Hz, using the g.USBamp amplifier (g.tec medical engineering GmbH).

**Figure 1.**
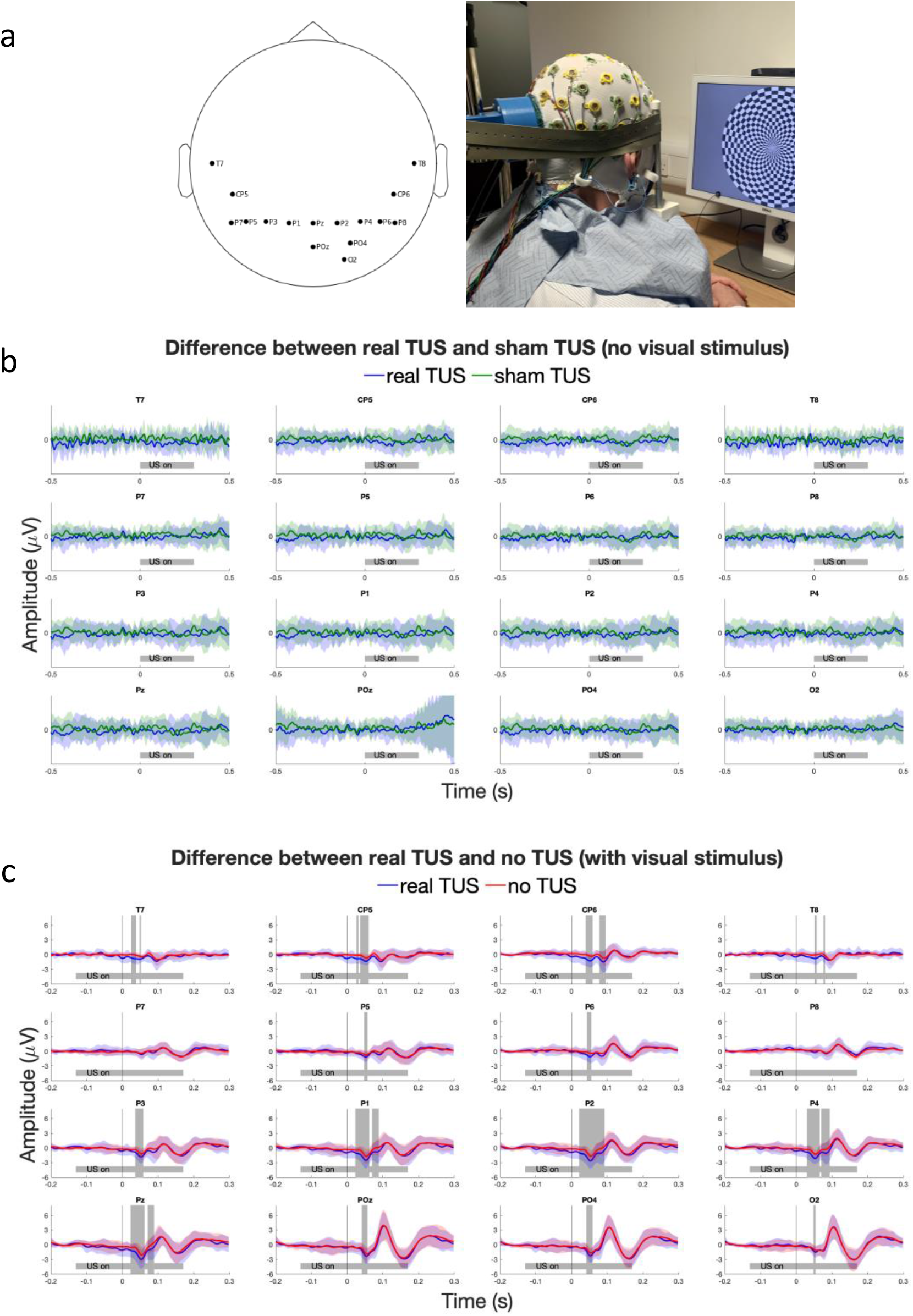
Depiction of recorded EEG channels and experimental set-up (a). EEG recordings in TUS only blocks (b), and TUS+checkerboard blocks (c) where vertical grey line represents checkerboard flip. No significant differences were found between sham TUS and no TUS in TUSonly blocks. Shaded grey areas in ‘c’ represent the windows in which significant differences were found between real TUS and no TUS conditions.

Participants were positioned in a chin rest. The transducer, connected to an articulated arm, was manually positioned 2 cm to the left of the inion and held in place using rubber straps. Acoustic coupling was achieved using a gel pad and ultrasound gel. Participants were dark adapted for 2 min before starting stimulation and then sat facing a screen 50-60 cm away. First, in two TUS-only blocks, real and sham pulses (100 each), were applied randomly at a fixed inter-stimulus interval of 2 s. Then, two TUS+checkerboard blocks were performed, where a checkerboard stimulus (checks at 25 and 75% of maximum screen luminance, and ∼32° visual angle) was flipped every 0.5 s, and every fourth stimulus was associated with either a real or sham (100 each) TUS trial. The TUS trial started 130 ms (± 0-5 ms jitter) before the checkerboard flip. Real and sham trials within each block were randomised and double-blind.

EEG data were analysed using eeglab (https://eeglab.org/) and FieldTrip (http://fieldtriptoolbox.org). The data were epoched relative to the TUS onset (−0.5 to 0.5 s) for TUS-only blocks, and relative to the checkerboard flip (−0.2 to 0.3 s) for TUS+checkerboard blocks. The following filters were applied: 50 Hz bandstop, 45 Hz lowpass and 1 Hz highpass, and the data were baseline corrected (−100 to 0 ms TUS-only, and -200 to -140 TUS+checkerboard). Across participants and conditions 14±7 trials were rejected using z-value based artifact detection with a cut-off of 15. After time-locked averaging, cluster-based permutation testing was used to examine differences between pairs of conditions across all channels.

In the TUS-only blocks, we did not observe any evoked potentials, and found no significant differences between real and sham TUS conditions (Fig. 1b). In contrast to previous reports, none of our participants reported phosphenes. As would be expected, in the TUS+checkerboard blocks, a clear VEP in response to the visual checkerboard was observed in all conditions. A statistically significant difference was observed between real and no TUS trials, in posterior electrodes, during a time period corresponding to the N75 component of the VEP, which likely originates in V1 (Fig. 1c). There was no difference between the sham and no TUS condition, but also no significant difference when the real and sham conditions were directly compared.

Our findings differ from previous work which showed that TUS applied to the V1 evokes a ‘VEP-like’ potential (1). While we used the same intensity as this previous study, in order to implement effective ramping, we lowered the PRF. Though some *in vitro* and animal data suggest that higher PRFs lead to stronger effects (6,7), the parameter space has not been extensively mapped, and in humans neuromodulatory effects have been demonstrated at PRFs similar to ours (8). As such, while the differences in parameters may have contributed to the lack of TUS-evoked activity in our study, we cannot rule out the possibility that effects reported previously were influenced by an auditory confound, in particular since the limited number of EEG channels did not allow localizing the anatomical origin of the evoked potential.

Adding to previous work showing modulatory TUS effects (2), we observed increased amplitude of the VEP N75 component. Modulation of the N75 amplitude has also been reported in response to transcranial magnetic and electrical stimulation (9), and can be accompanied by a change in contrast sensitivity (10). Our data suggest that TUS can be added to the repertoire of non-invasive brain stimulation tools to study the physiological basis of visual perception, and more generally, to modulate neural activity for basic science and clinical applications. However, we did not find any differences between sham and real TUS conditions, likely due to fewer trials in these conditions compared to the no TUS VEPs (100 vs 300), but replication studies are required to draw definitive conclusions.

In conclusion, we demonstrate that ramped TUS, which is less audible than pulsing regimes with sharp onsets and offsets, can modulate neural activity. This suggests that, in line with in-vitro and animal data, there is a direct neuromodulatory effect of ultrasound, in addition to any confounding effects. Moving forward, ramping offers a relatively easy approach to minimise the auditory confound.

## Acknowledgements

This work was supported by the Engineering and Physical Sciences Research Council UK (EP/P008860/1, EP/P008712/1, EP/S026371/1). EM was supported by a UKRI Future Leaders Fellowship (MR/T019166/1) and in part by the Wellcome/EPSRC Centre for Interventional and Surgical Sciences (WEISS) (203145Z/16/Z). SB was supported by the Dunhill Medical Trust, (RPGF1810\93). TN was supported by a grant from the Boehringer Ingelheim Foundation (BIS) to TOB. The research was supported by the National Institute for Health Research (NIHR) Oxford Biomedical Research Centre and the NIHR Oxford Health Biomedical Research Centre. The Wellcome Centre for Integrative Neuroimaging is supported by core funding from the Wellcome Trust (203139/Z/16/Z). CJS holds a Senior Research Fellowship, funded by the Wellcome Trust (224430/Z/21/Z).

